# A repressive regulatory cascade shapes temporal patterning of activity-regulated gene expression in a defined sensory neuron type

**DOI:** 10.64898/2026.05.14.725236

**Authors:** Samuel G. Bates, Nathan Harris, Stephen Nurrish, Piali Sengupta

## Abstract

Long-term neuronal plasticity is driven by activity-regulated gene (ARG) expression programs that encode stimulus features in a neuron type-specific manner ^1–6^. ARG programs are typically characterized by rapid induction of immediate early genes (IEGs) without requiring new protein synthesis, followed by expression of secondary response genes regulated by IEG-encoded transcription factors ^2,5,7–14^. However, the molecular mechanisms that pattern these programs in specific neuron types *in vivo* in response to physiological stimuli are not fully described. We previously showed that temperature regulates an ARG program in the *C. elegans* AFD thermosensory neuron pair to drive behavioral plasticity ^3,15,16^. By profiling AFD following temperature upshifts of varying duration, here we show that ARGs in this neuron exhibit distinct temporal trajectories. Notably, rapidly induced genes do not include known IEGs but are enriched for molecules implicated in signal transduction and navigation. Both rapid and delayed ARG expression require the CMK-1 CaMKI kinase and CRH-1 CREB transcription factor, with CRH-1 acting at both early and late stages. We further define a temporal regulatory cascade in which CREB-dependent rapid induction of the RCAN-1 calcineurin regulator acts in parallel with the MEF-2 transcription factor to repress premature expression of a delayed ARG. Subsequent downregulation of RCAN-1 likely enables CRH-1-dependent ARG expression at later stages. Our results demonstrate that in addition to classical gene-activating transcriptional cascades, ARG-controlled repressive mechanisms also operate to precisely shape the temporal dynamics of an ARG cascade in a sensory neuron type *in vivo*, and suggest that distinct cell type-specific regulatory pathways operate to drive ARG expression programs across neuron types.

## RESULTS and DISCUSSION

### The duration of temperature experience is encoded in temporally regulated gene co-expression modules in the AFD thermosensory neuron pair

To determine how the duration of temperature experience is reflected in the molecular profile of AFD, we shifted animals grown at 15°C overnight to 25°C for 1 hr or 4 hr, and isolated ribosome-associated RNA from the two AFD neurons via translating ribosome affinity purification followed by sequencing (TRAP-seq) ^15,17^ (Figure 1A). Differential expression analyses against whole animal lysates showed enrichment of known markers of AFD identity and function in the AFD samples (Figure S1A), confirming successful isolation of AFD-specific transcripts.

**Figure 1.**
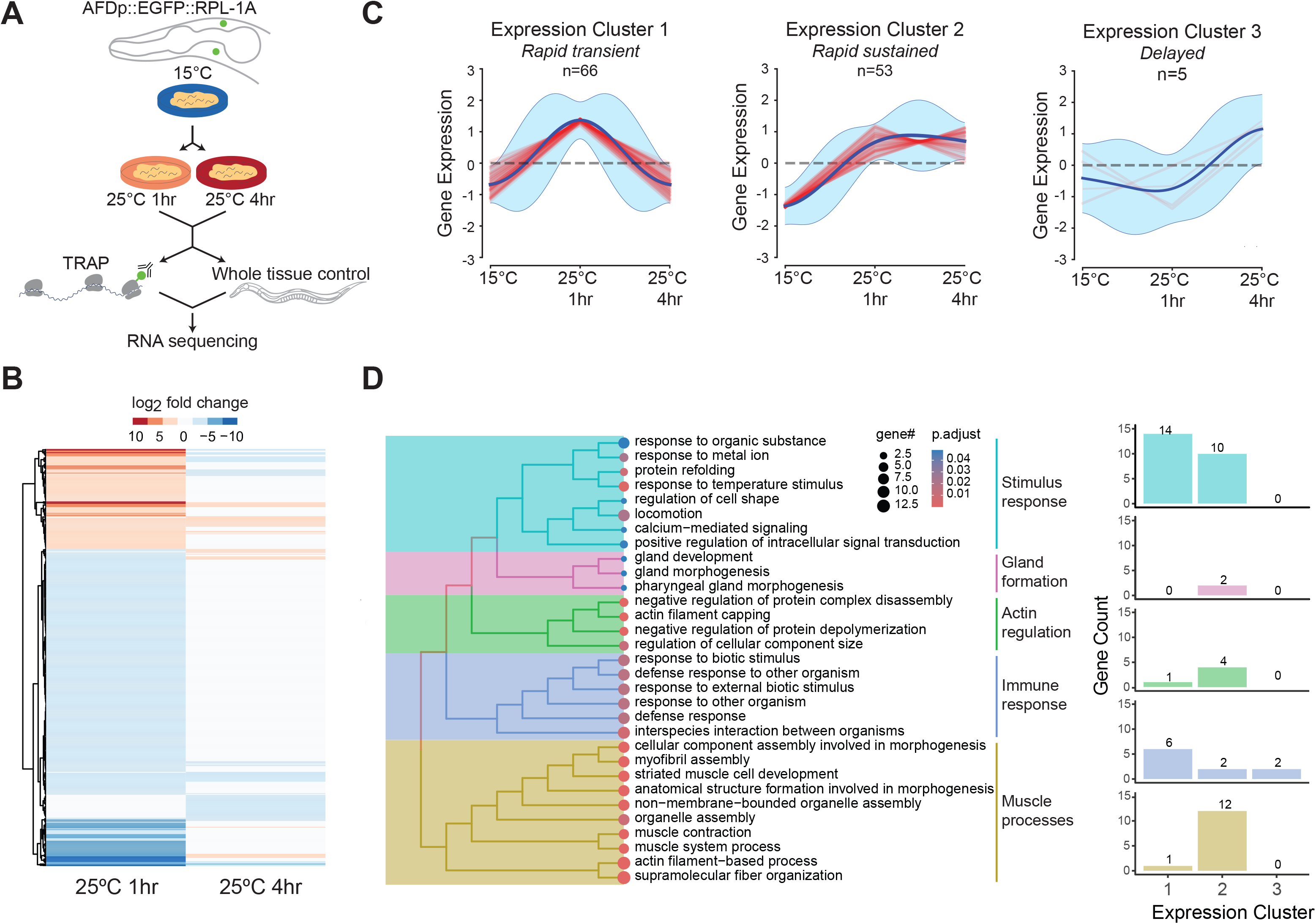
Identification of temperature-regulated gene co-expression modules in AFD via TRAP-seq. **A)** Schematic of experimental design to identify temperature-regulated transcripts in AFD via TRAP-seq. **B)** Heatmap showing differentially expressed genes in AFD following a temperature upshift to 25°C for 1 hr or 4 hr. Shown genes are filtered for ≥ log2-fold change and p value ≤ 0.05. Data are from 4 independent TRAP-seq experiments. **C)** Grouping of genes upregulated upon a temperature upshift as shown in B into co-expression clusters ^21^. Red lines: expression trajectories of individual genes; dark blue lines: mean expression; light blue shading: ± 2X SD. **D)** (Left) GO terms enriched in the upregulated gene set summarized using hierarchical clustering by semantic similarity. (Right) Number of genes in each expression cluster categorized by GO terms shown at left. Also see Figure S1.

We next identified genes whose expression trajectories in AFD are altered by temperature experience relative to whole animal controls. At 1 hr, the majority of differentially expressed genes (∼60%) was downregulated (Figure 1B); the expression of a large fraction of these genes returned to baseline at 4 hr (Figure 1B). Since we and others previously described the contributions of several upregulated genes to AFD functional plasticity (eg. ^3,15,18–20^), we focused subsequent analyses on upregulated genes; downregulated genes will be described elsewhere.

Upregulated genes were clustered into co-expression modules using an infinite Gaussian process mixture algorithm (DPGP: Dirichlet process Gaussian process mixture model) that groups genes with similar temporal expression profiles from time-series data ^21^. This analysis identified three distinct expression trajectories (Figure 1C). The largest cluster showed rapid but transient induction returning to baseline by 4 hr (expression cluster 1: *rapid transient*; Figure 1C). A second cluster was also rapidly induced but levels remained elevated at 4 hr (expression cluster 2: *rapid sustained*; Figure 1C). The smallest cluster exhibited delayed induction, with significant upregulation only at 4 hr (expression cluster 3: *delayed*; Figure 1C). Similar rapid transient and rapid sustained but not delayed patterns were also observed in whole animal samples, with an additional slow-rising cluster (*slow*; Figure S1B). Together, these results extend previous analyses of a limited gene set in AFD ^15^ and establish that ARGs in AFD form temporally distinct modules that reflect the duration of temperature experience.

Time-series profiling of ARG programs suggest that IEGs include but are not restricted to molecules that are partly shared across different neuron types, whereas neuron type-specific effector genes are induced in subsequent waves ^2,6,14,22,23^. To determine whether a similar pattern is also observed in the temperature-regulated expression program in AFD, we performed gene ontology analyses on the set of upregulated genes. We found that genes implicated in signaling including in temperature responses and taxis behaviors, were enriched in this set and distributed across clusters 1 and 2 (Figure 1D, Figure S1C), indicating that these genes are rapidly upregulated upon a temperature upshift. Genes assigned to enriched terms related to actin-myosin regulation were predominantly members of Cluster 2 (Figure 1D, Figure S1C). Temperature experience modulates the morphology of the complex actin-based microvilli comprising the AFD sensory endings ^24–26^; these molecules may mediate activity-dependent modification of AFD sensory ending architecture. In contrast, genes upregulated in whole animal samples were distributed across multiple gene categories (Figure S1C), consistent with general temperature-but not activity-mediated regulation. Notably, rapidly induced genes in AFD did not include canonical IEGs such as *fos-1, jun-1*, and *egr1*-related genes identified in ARG programs across cell types in multiple organisms ^27,28^. These results indicate that temperature experience rapidly induces genes in AFD predicted to regulate neuronal responses and behavior.

### Transcriptional mechanisms contribute to temperature-regulated gene expression changes

TRAP-Seq measures association of mRNAs with ribosomes; thus, altered mRNA levels identified via this method may reflect both transcriptional and non-transcriptional regulation. We previously showed that experience-dependent upregulation of a subset of ARGs in AFD is mediated in part by increased transcription ^15,16^. To validate these expression changes and define the required regulatory mechanisms, we characterized the expression of representative ARGs from each upregulated co-expression cluster.

To assess transcriptional activity in specific cell types *in vivo,* we inserted an *SL2* trans-splice leader sequence followed by reporter sequences into the 3’UTRs of endogenous loci via genome editing ^29^. Since a subset of candidate genes is expressed in cells other than AFD, we inserted *SL2::H2B::gfp11* ^30^ sequences into the endogenous loci and expressed *gfp(1-10)* as a single copy transgene under a temperature-insensitive AFD-specific promoter (see Methods) to enable visualization of reconstituted split-GFP specifically in AFD.

A subset of genes in the rapid transient cluster exhibited expression dynamics consistent with both transcriptional and post-transcriptional regulation. For example, the *gcy-18* receptor guanylyl cyclase thermoreceptor gene ^20^ exhibited rapid transient changes in TRAP-seq analysis (Figure 2A) similar to the previously characterized rapidly upregulated *pyt-1* adaptor gene ^15^. However, analyses of both mRNA levels in AFD via qRT-PCR ^16^ and quantification of *gcy-18::SL2::gfp* reporter expression indicated that while *gcy-18* transcription is rapidly induced, the expression of this gene is subsequently maintained upon prolonged exposure to warmer temperatures (Figure 2B, 2C). Consistently, endogenously tagged GCY-18::GFP fusion protein levels are increased at the AFD sensory endings ∼4 hr after the temperature upshift and are then maintained, in part via increased dendritic trafficking ^3,15^. Together, these results indicate that multiple mechanisms contribute to modulating functional GCY-18 protein levels in AFD upon warming.

**Figure 2.**
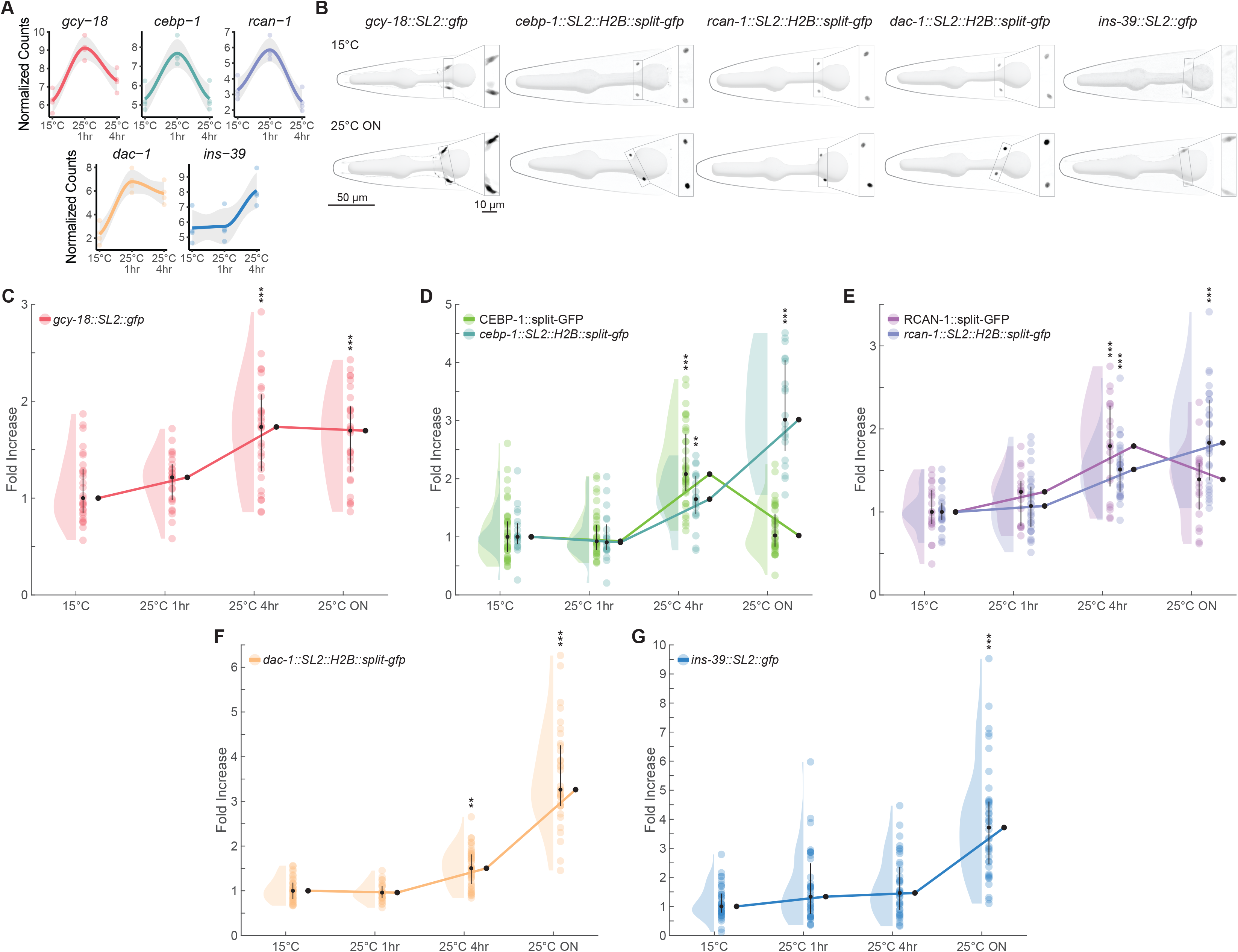
Temperature-regulated gene induction in AFD is mediated in part via transcriptional mechanisms. **A)** Gene expression trajectories measured by TRAP-seq. Circles are gene-wise counts normalized by DESeq2. Lines are functions fit to circles by Loess regression. Shaded regions represent confidence intervals. **B)** Representative images showing expression of the indicated reporter genes in the two AFD neurons in the head of an adult hermaphrodite. Insets indicate AFD soma. Anterior is at left. **C-G)** Quantification of fold-increase of fluorescence intensities of the indicated transcriptional and translational reporters in AFD at 15°C, and at the shown timepoints following a shift from 15°C to 25°C. Each circle is a measurement from a single AFD soma normalized to the median at 15°C. Black circles and vertical lines show median and quartiles, respectively. ** and ***: different at P<0.01 and 0.001, respectively compared to values at 15°C (one-way ANOVA with Dunnett’s correction). ≥2 independent experiments each.

The *cebp-1* C/EBP transcription factor, and the *rcan-1* RCAN1 calcineurin regulator genes also exhibited rapid transient expression changes in TRAP-seq data (Figure 2A). Both endogenously tagged CEBP-1 and RCAN-1 fusion proteins and *SL2::H2B::gfp11* transcriptional reporters were rapidly induced (Figure 2D, 2E), indicating regulation via transcriptional mechanisms. However, while levels of both proteins decreased after overnight exposure to 25°C, both transcriptional reporters showed further induction (Figure 2B, 2D, 2E). These observations suggest that these proteins may be downregulated at late timepoints via post-transcriptional and/or post-translational mechanisms. RCAN1 and C/EBP proteins have previously been shown to be degraded by ubiquitination and proteosome-mediated degradation as well as chaperone-mediated autophagy ^31–34^.

The *dac-1* Dachsund transcription factor ^15,35^ exhibited rapid sustained upregulation in AFD as measured using an endogenously tagged *SL2::H2B::gfp11* reporter (Figure 2A, 2B, 2F), consistent with the expression dynamics of the endogenously tagged *dac-1::gfp* fusion protein ^15^. Finally, significant induction of the *ins-39* insulin-like peptide transcriptional reporter ^15,19,36^ was observed only following prolonged exposure to 25°C (Figure 2B, 2G) ^15^, consistent with the assignment of this gene to the delayed upregulated cluster (Figure 2A). The delay in the timing of reporter induction as compared to the TRAP-seq trajectories likely reflect fluorophore maturation kinetics, and/or other regulatory pathways.

Taken together, these data indicate that temperature experience-dependent expression changes in AFD are primarily mediated via transcriptional regulation, but that post-transcriptional and/or post-translational mechanisms likely further shape their temporal expression trajectories.

### CRH-1 CREB acts both early and late to regulate gene expression changes upon a temperature upshift

Calcium influx upon neuronal depolarization activates transcription factors such as CREB, SRF1, CaRF and MEF2 through CaMKs and MAPK/ERK signaling pathways, thereby coupling neuronal activity to gene expression programs ^2,37–41^. We previously showed that rapid upregulation of the *pyt-1* adaptor molecule requires both CMK-1 CaMK I/IV and CRH-1 CREB following temperature upshift, whereas *dac-1* induction is CMK-1-dependent after overnight warming but not at 4 hr, and is only partially CRH-1-dependent at both timepoints ^15^. Temperature-dependent induction of other rapidly upregulated genes is CRH-1-independent ^15^, indicating that multiple activity-regulated pathways induce gene expression in AFD. We tested whether transcriptional upregulation of subsets of ARGs identified in this work is dependent on CMK-1 and CRH-1.

Induction of both *rcan-1* and *cebp-1* transcriptional reporters was partly or fully dependent on CRH-1 and CMK-1 at 4 hr after temperature upshift (Figure 3A, 3B). At the overnight timepoint, induction of *cebp-1* but not *rcan-1* was partly CMK-1-dependent (Figure 3A, 3B). *rcan-1* expression levels were decreased at 15°C in *crh-1* mutants, whereas *cebp-1* levels were increased under these conditions (Figure 3A, 3B). In contrast, mutations in either *crh-1* or *cmk-1* had little effect on *gcy-18* upregulation (Figure 3C). *gcy-18* expression was strongly reduced at 15°C in *cmk-1* mutants (Figure 3C); these low expression levels may have precluded detection of further upregulation in *cmk-1* mutants in qRT-PCR assays ^16^. Mutating a high confidence CREB binding motif (CRE: cAMP response element) 132 bp upstream of the *gcy-18* initiator ATG at the endogenous locus abolished *gcy-18* induction (Figure S2A), suggesting that an ATF/CREB transcription factor other than CRH-1 may mediate *gcy-18* upregulation.

**Figure 3.**
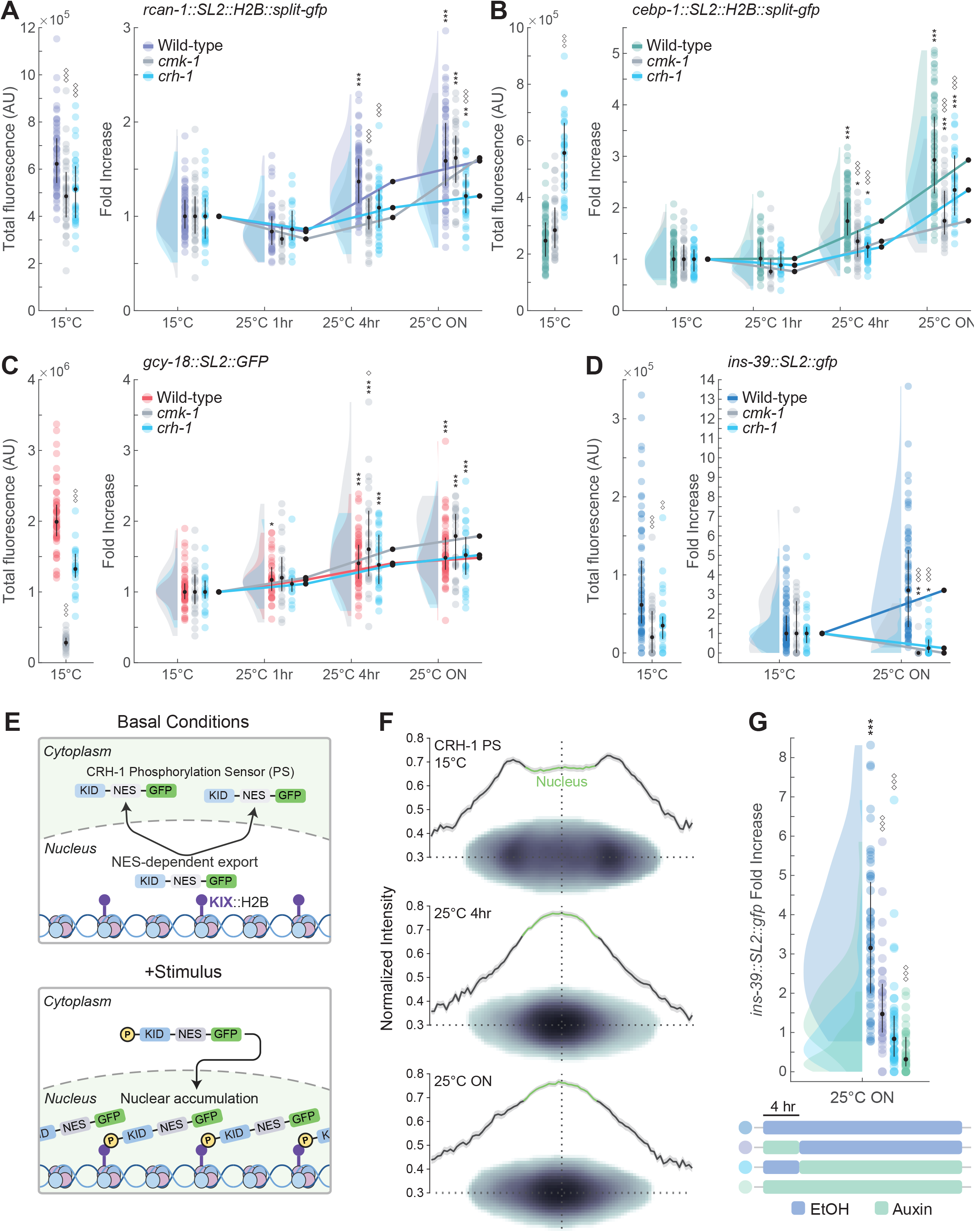
CRH-1 CREB activity is necessary at early and late timepoints to induce activity-regulated gene expression. A-D) (Left) Quantification of fluorescence intensities of the indicated transcriptional reporters in AFD at 15°C in the shown genetic backgrounds. Each circle is a measurement from a single AFD soma. (Right) Quantification of fold-increase of fluorescence intensities of transcriptional reporters in the indicated temperatures and temperature shift conditions in the shown genetic backgrounds. Each circle is a measurement from a single AFD soma normalized to the median at 15°C. Black circles and vertical lines show median and quartiles, respectively. *, ** and ***: different at P<0.05, 0.01 and 0.001, respectively compared to values at 15°C within each genotype (one-way ANOVA with Dunnett’s correction). ◊, ◊◊◊: different at P<0.05 and 0.001, respectively compared to wild-type values within a temperature condition (one-way ANOVA with Dunnett’s correction). 3 independent experiments (A-C), 2 independent experiments (D). Alleles used were *cmk-1(oy21)* and *crh-1(tz2)*. **E)** Cartoon of CRH-1 phosphorylation sensor (CRH-1 PS) design. In the basal state, KID::GFP resides primarily in the cytoplasm of AFD due to the inclusion of nuclear export sequences (NES). Upon activation, phosphorylation of KID promotes interaction with KIX::H2B and increases nuclear GFP accumulation in AFD. **F)** CRH-1 phosphorylation sensor (CRH-1 PS) subcellular signal distribution at the indicated temperature conditions as shown by mean projections of nucleus-aligned images and mean line scan profile of all aligned images. Shaded area around the mean line scan is +/-SEM. n>150 AFD cells per condition; 8 independent experiments. **G)** (Top) Fold-increase in *ins-39::SL2::gfp* fluorescence intensity normalized to 15°C in the conditions shown schematically (bottom). Each circle is a measurement from a single AFD soma normalized to the median at 15°C. Black circles and vertical lines show median and quartiles, respectively. ***: P<0.001 compared to values at 15°C within each condition (one-way ANOVA with Dunnett’s correction). ◊◊◊: different at P<0.001 compared to the no auxin control (one-way ANOVA with Dunnett’s correction). 4 independent experiments. Also see Figure S2.

We next tested whether the delayed induction of *ins-39* following a temperature upshift is also CMK-1 and/or CRH-1-dependent. Although *ins-39* expression was also reduced at 15°C in *cmk-1* and *crh-1* mutants, induction of this gene upon prolonged warming was fully dependent on both CRH-1 and CMK-1 (Figure 3D). Deletion of a single predicted CRE 34 bp upstream of the initiator ATG in *ins-39* upstream regulatory sequences had no effect on induction of expression (Figure S2B) suggesting that additional regulatory elements and/or indirect CRH-1-dependent mechanisms contribute to the induction of *ins-39* ^36^. We conclude that both rapid and delayed induction of a subset of genes in AFD is regulated by a CMK-1/CRH-1 cascade, but that additional mechanisms also operate to alter gene expression in response to a temperature upshift. The deployment of multiple regulatory pathways to alter the expression of different gene subsets may enable integration of distinct stimulus features and fine-tuning of gene expression dynamics ^42,43^.

Upon a temperature upshift, AFD exhibits a phasic increase in intracellular calcium levels which returns to baseline within minutes despite continued exposure to the warmer temperature ^44^. A role for CRH-1 in regulating *ins-39* expression ∼24 hr post temperature upshift was unexpected given the well-characterized role for this transcription factor in the induction of IEGs ^45,46^ and a subset of rapidly upregulated genes in AFD ^15^ (Figure 3A, 3B). To determine whether CRH-1 activity is transient or persists upon prolonged warming, we first designed an *in vivo* sensor to monitor CRH-1 phosphorylation in AFD. CRH-1 is activated upon phosphorylation at the S48 residue (equivalent to S133 in mammalian CREB) in its kinase-inducible domain (KID), enabling interaction with the KID-interacting domain (KIX) of the CBP-1 CREB coactivator ^47–49^. Thus, the KID-KIX interaction provides a readout of the phosphorylation state of CRH-1 ^50–52^. We fused the CRH-1 KID to a nuclear export sequence and GFP and expressed this fusion protein together with CBP-1 KIX fused to histone H2B in AFD (Figure 3E). In the absence of phosphorylation, GFP is enriched in the cytoplasm, whereas following phosphorylation, the KID-KIX interaction retains GFP in the nucleus (Figure 3E). Subsequent dephosphorylation would allow GFP to be exported back to the cytoplasm. GFP from the CRH-1 phosphorylation sensor (CRH-1 PS) was predominantly cytoplasmic at 15°C, but became significantly enriched in the nucleus upon a shift to 25°C for 4 hr consistent with CRH-1 activation (Figure 3F, Figure S2C, S2D). Nuclear enrichment of the sensor persisted after overnight exposure to 25°C (Figure 3F, Figure S2C, S2D), indicating that CRH-1 remains phosphorylated for a prolonged period following initial activation. An S48A mutation in KID significantly decreased nuclear GFP accumulation under all temperature conditions (Figure S2C, S2E), confirming that phosphorylation of S48 drives sensor localization.

We next tested whether CRH-1 function is also necessary at later timepoints following warming to regulate gene expression. To spatiotemporally deplete CRH-1 in AFD we tagged *crh-1* at its endogenous locus with degron sequences, expressed the auxin receptor TIR1 specifically in AFD as a single copy transgene.^53,54^, and exposed animals to auxin at different times during a temperature upshift. Auxin was present throughout the 24 hr time course, only during the first 4 hr, or only during the final ∼20 hr after temperature upshift (Figure 3G). Continuous CRH-1 depletion or depletion only during later stages abolished *ins-39* induction (Figure 3G), indicating that CRH-1 is required cell-autonomously at later timepoints to upregulate *ins-39* expression. Depleting CRH-1 during the first 4 hr following the temperature upshift also significantly reduced *ins-39* induction (Figure 3G) although we are unable to exclude the possibility that CRH-1 levels were not fully restored following removal of auxin. We infer that CRH-1 activity is required at both early and late phases of the temperature response in AFD to induce rapid and delayed gene expression.

### Delayed upregulation of *ins-39* expression is mediated by temporal control of a repressive pathway

Although *ins-39* expression is both CMK-1 and CRH-1-dependent similar to the regulation of subsets of rapidly induced genes, upregulation of *ins-39* is observed only following prolonged exposure to warm temperatures. Delayed induction of secondary response genes following neuronal stimulation has been shown to be mediated by IEG transcription factors ^6,11,12,27,39^. The CEBP-1 transcription factor is induced rapidly (Figure 2D), but mutations in *cebp-1* only partly downregulated *ins-39* expression (Figure S3A).

An alternative mechanism regulating delayed expression is the transient induction of a repressive molecule that antagonizes activation at early timepoints. RCAN proteins regulate gene expression primarily via modulation of the calcium/calmodulin-dependent phosphatase calcineurin depending on expression levels and cellular context ^55–60^, thereby modulating the phosphorylation state and activity of downstream transcription factors and co-factors ^56,61–63^. We tested the hypothesis that rapid induction of RCAN-1 (Figure 2E) antagonizes *ins-39* expression, thereby delaying its upregulation. *ins-39* expression was increased significantly although to a minor extent at 4 hrs after temperature upshift in *rcan-1* mutants (Figure 4A, 4B, Figure S3B). Since neuronal activity regulates gene expression via multiple pathways ^13,39^, we considered the possibility that *ins-39* expression is repressed by mechanisms acting in parallel with RCAN-1. The MEF-2 transcription factor has been implicated in activity-regulated gene expression across different cell types and species ^64–70^, and can function as either an activator or repressor based on its phosphorylation state and its association with co-regulatory proteins ^71,72^. Although *mef-2* mutants alone did not exhibit altered *ins-39* expression at the 4 hr timepoint, *ins-39* expression was significantly upregulated in *mef-2; rcan-1* double mutants at 4 hr (Figure 4A, 4B, Figure S3B). Expression levels in this double mutant were also elevated and highly variable at 15°C (Figure 4B, Figure S3B, S3C). Increased *ins-39* expression in the *mef-2; rcan-1* double mutants was abolished in *crh-1* mutants at all timepoints (Figure 4B, Figure S3B). Consistent with increased activation of CRH-1 in *rcan-1* mutants, CRH-1 phosphorylation was enhanced in this mutant background (Figure 4C, 4D, Figure S3D). We conclude that RCAN-1 and MEF-2 act in parallel to prevent CRH-1-mediated induction of *ins-39* expression both under basal conditions and at early timepoints following a temperature upshift.

**Figure 4.**
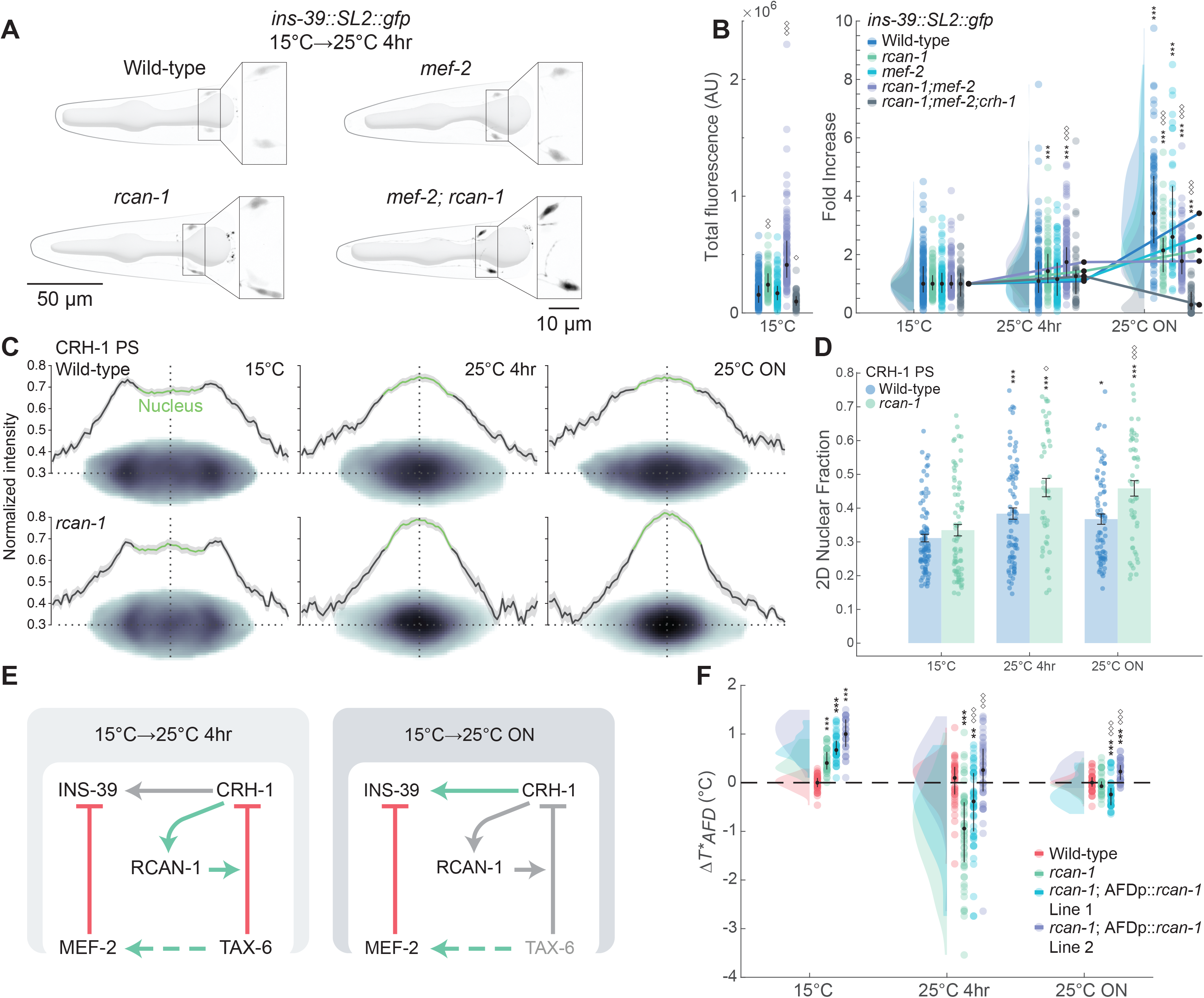
Transient induction of an RCAN-1-dependent repressive pathway prevents premature expression of *ins-39*. **A)** (Top) Representative images showing expression of *ins-39::SL2::gfp* in the two AFD neurons in the head of an adult hermaphrodite in the indicated genetic backgrounds following a 4 hr temperature upshift. Insets indicate AFD soma. Anterior is at left. **B)** (Left) Quantification of fluorescence intensities of the *ins-39* transcriptional reporter in AFD at 15°C in the shown genetic backgrounds. Each circle is a measurement from a single AFD soma. (Right) Quantification of fold-increase of fluorescence intensities of the *ins-39* transcriptional reporters in the indicated temperatures and temperature shift conditions in the shown genetic backgrounds. Each circle is a measurement from a single AFD soma normalized within day to the median at 15°C. Black circles and vertical lines show median and quartiles, respectively. ***: different at P<0.001 compared to values at 15°C within each genotype (one-way ANOVA with Dunnett’s correction). ◊, ◊◊, ◊◊◊: different at P<0.05, 0.01 and 0.001, respectively, from wild-type within a temperature condition (one-way ANOVA with Dunnett’s correction). 10 independent experiments. Alleles used were *mef-2(gv1), rcan-1(tm1925),* and *crh-1(tz2)*. **C)** CRH-1 PS subcellular signal distribution in AFD in wild-type (top) and *rcan*-*1(tm1925)* mutant animals (bottom) in the indicated temperature conditions as shown by mean projections of nucleus-aligned images and mean line scan profiles of all aligned images. Shaded area around the mean line is +/-SEM. n>40 AFD cells per genotype/condition; 4 independent experiments. **D)** Quantification of CRH-1 PS nuclear signal intensity in AFD in the shown temperature conditions. Values are calculated from individual mean projections as the sum nuclear intensity divided by total cell intensity. Each circle is the value from a single AFD neuron. Horizontal and vertical bars indicate mean and SEM, respectively. *, ** and ***: different at P<0.05, 0.01 and 0.001, respectively, compared to values at 15°C within each genotype (one-way ANOVA with Dunnett’s correction). ◊, ◊◊◊: different at P<0.05 and 0.001, respectively, from wild-type within a temperature condition (one-way ANOVA with Dunnett’s correction). Images and quantification of wild-type samples are a subset of those shown in Figure 3F and Figure S2C, respectively. 4 independent experiments. **E)** Proposed working model for the pathways regulating the temporal dynamics of activity-regulated *ins-39* expression in AFD. At 4 hrs after temperature upshift*, rcan-1* is induced in a CRH-1-dependent manner. RCAN-1 subsequently acts through TAX-6 calcineurin or potentially additional pathways to inhibit CRH-1 activity and repress *ins-39* expression. In parallel, MEF-2 antagonizes CRH-1-dependent transcription. In *rcan-1* mutants, TAX-6-dependent inhibition of CRH-1 is alleviated, but MEF-2 repression persists resulting in a dampening of *ins-39* induction. Conversely, in *mef-2* mutants, RCAN-1-dependent repression of CRH-1 activity through TAX-6 is maintained. Loss of both *rcan-1* and *mef-2* together enables premature CRH-1-dependent *ins-39* expression. At later timepoints, downregulation of RCAN-1 contributes to the release of repression, thereby permitting CRH-1-mediated induction of *ins-39*. **F)** *T*_AFD_* in animals of the indicated genotypes relative to wild-type at each temperature condition. The mean of all wild-type animals is set as the baseline at each temperature condition (horizontal dotted line). For each data point, the wild-type mean was subtracted to calculate the difference between that data point and the wild-type mean within each condition. Wild-type *rcan-1* sequences were expressed in AFD under the *gcy-8* promoter as a stably integrated single copy transgene. ** and ***: different at P<0.01 and 0.001, respectively compared to wild-type within a temperature condition (one-way ANOVA with Dunnett’s correction). ◊◊◊: different at P<0.001 from *rcan-1* mutants within a temperature condition (one-way ANOVA with Dunnett’s correction). 3 independent experiments. The *rcan-1(tm1925)* allele was used. Also see Figures S3 and S4.

Calcineurin regulates gene expression by targeting multiple transcription factors including CREB and MEF2 ^62,71,73–78^. In *C. elegans*, the TAX-6 calcineurin homolog has previously been implicated in regulating AFD thermosensory function ^79,80^. To test whether calcineurin activity contributes to *ins-39* regulation, we examined *ins-39* expression in *tax-6(gof)* and *tax-6(lof)* mutants (Figure S3E). Although *ins-39* expression at the 4 hr timepoint was unaffected, induction at the overnight timepoint was abolished in *tax-6(gof)* mutants (Figure S3E). Sustained synaptic input has been shown to result in the persistence of phosphorylated CREB in part via inactivation of calcineurin-dependent inhibition of CREB dephosphorylation ^73,74,81,82^. Since the *tax-6(gof)* allele encodes a predicted constitutively active calcium-independent enzyme ^83^, persistent phosphatase activity may prevent induction of *ins-39* expression by constitutive dephosphorylation of CRH-1. In wildtype animals, downregulation of RCAN-1 at later timepoints and/or loss of calcium-dependent regulation of calcineurin activity may relieve TAX-6-dependent inhibition of CRH-1 and permit *ins-39* induction. Consistently, in the absence of *tax-6* in *tax-6(lof)* mutants, *ins-39* expression was upregulated at 15°C and did not further increase upon a temperature upshift (Figure S3E). We hypothesize that RCAN-1 acts in part via TAX-6 calcineurin to shape *ins-39* induction dynamics.

Together, these data support a model in which neuronal activity induces *rcan-1* transcription via CRH-1 following temperature upshift. Elevated RCAN-1 then feeds back through TAX-6 calcineurin or potentially additional pathways ^84^ to repress CRH-1-dependent *ins-39* expression (Figure 4E). MEF-2 antagonizes CRH-1-dependent *ins-39* upregulation at the 4 hr timepoint possibly via its calcineurin-regulated phosphorylation state and/or interaction with HDACs ^68,85–87^ but does not itself promote expression. MEF-2 may repress *ins-39* induction via competition with CRH-1 for recruitment of co-regulators or modulation of chromatin state (eg. ^88,89^). For example, the NCoR repressor complex is pre-bound with CREB at the promoters of rapidly upregulated genes in cultured primary neurons and prevents premature CREB-dependent upregulation despite their open chromatin state ^2^. In the absence of both *rcan-1* and *mef-2* activity, *ins-39* expression is upregulated prematurely via CRH-1. At later timepoints, reduced RCAN-1 protein levels (Figure 2E) contribute to the relief of this inhibition, thereby enabling CRH-1-dependent induction of *ins-39* (Figure 4E). INS-39 plays an important role in regulating hermaphrodite survival under stress conditions ^19,36^, and in addition to temperature, the expression of this gene is regulated by sex and developmental stage ^19,36^. These observations suggest that multiple inputs are integrated at the level of transcription to precisely regulate *ins-39* expression dynamics.

### Mutations in *rcan-1* alter AFD thermosensory response plasticity

The expression dynamics of *rcan-1* likely contributes to shaping the temporal dynamics of genes other than *ins-39* in AFD, possibly including genes that regulate experience-dependent plasticity in AFD responses. The response threshold of AFD (*T*_AFD_*) shifts to a higher temperature on a minutes and hours-long timescale upon exposure to warmer temperatures ^16,44,90–93^. *T*_AFD_* changes on longer timescales require a temporally regulated ARG expression program in AFD ^15,16,94^. As an example, we previously showed that the rapid transient upregulation of the *pyt-1* adaptor molecule following AFD activation is required for precise temporal tuning of *T*_AFD_* only during the time period during which *pyt-1* is induced ^3,15^.

Previous work reported that *T*_AFD_* is not altered in *rcan-1* mutants grown at either 15°C or 25°C overnight ^80^. However, whether *rcan-1* contributes to AFD response plasticity following an acute temperature change was not examined. We found that *rcan-1* mutants exhibited significantly lower *T*_AFD_* at 4 hr after the temperature upshift (Figure 4F, Figure S4). This defect was rescued upon AFD-specific expression of *rcan-1* (Figure 4F, Figure S4). Conversely, *T*_AFD_* shifted to warmer temperatures in *rcan-1* mutants after overnight growth at 15°C, and these values were further increased upon constitutive expression of wild-type *rcan-1* sequences (Figure 4F, Figure S4). The discrepancy with prior results ^80^ may arise from the use of different calcium indicators or temperature conditions. These observations support the hypothesis that temporal regulation of RCAN-1 contributes to tuning AFD response plasticity as a function of temperature experience.

In summary, we describe an ARG expression program in the single AFD thermosensory neuron pair in *C. elegans* that encodes the duration of temperature change experienced by the animal. We identify ARG subsets whose expression is altered on distinct timescales, such that the gene expression profile of this neuron type at any timepoint provides a snapshot of the animal’s temperature history. Via temporal depletion of CRH-1 and analysis of CRH-1 phosphorylation state, we establish that CRH-1 is active and is required at both early and late timepoints for rapid and delayed gene expression. We also identify a novel regulatory strategy in which temporally regulated signaling and feedback repressive mechanisms shape the transcriptional dynamics in AFD.

Encoding a neuron’s activity history in its transcriptional program provides a mechanism for fine-tuning cellular and network properties over time. Sequential cascades of activity-regulated transcription factors are well-established mechanisms for temporally ordered gene expression programs ^27,40,42^. However, although IEGs including transcription factor genes are proposed to be broadly shared across neuron types as an early signature of neuronal activity, single cell profiling studies indicate that IEGs differ as a function of both neuronal identity and stimulus properties ^5,14,23,95,96^. In sensory systems, rapid induction of cell-specific signaling molecules may provide a particularly efficient mechanism to translate a defined sensory experience into adaptive changes in neuronal state, circuit function, and behavior ^4,15^. The deployment of distinct cell-specific IEG modules suggests that the regulatory architecture governing subsequent waves of gene expression may also differ across neuron types ^6,14^. Our identification of such an alternative strategy in AFD supports this hypothesis. Defining the diversity of ARG programs and their underlying regulatory logic will require comparisons of these pathways within and across defined neuron types *in vivo* in response to biologically relevant stimuli.

## SUPPLEMENTAL FIGURE LEGENDS

**Figure S1. Related to Figure 1. Characterization of temperature-regulated gene co-expression modules in whole animals.**

**A)** Enrichment of validated AFD-specific or-selective genes as measured by TRAP-Seq from AFD vs whole animal lysates as compared to single cell RNA-Seq data ^97^.

**B)** Co-expression clusters of upregulated genes in whole animals upon a temperature upshift. Red lines: expression trajectories of individual genes; dark blue lines: mean expression; light blue shading: ± 2X SD. Shown genes are filtered for ≥ 2-fold upregulation with p value ≤ 0.05 in whole animal but not AFD samples.

**C)** (Left) Fraction of unannotated genes in AFD-specific and whole animal expression clusters. (Right) GO term categorization of upregulated genes in AFD-specific and whole animal expression clusters summarized using hierarchical clustering by semantic similarity. Numbers indicate the number of genes in each expression cluster categorized by GO terms shown at left.

**Figure S2. Related to Figure 3. CREB activity persists to induce delayed *ins-39* expression A,B)** (Left) Quantification of fluorescence intensities of the indicated transcriptional reporters in AFD at 15°C in the shown genetic backgrounds. Each circle is a measurement from a single AFD soma. (Right) Quantification of fold-increase of fluorescence intensities of transcriptional reporters in the indicated temperatures and temperature shift conditions in the shown genetic backgrounds. CREmut: mutation of a single CREB-binding motif at the endogenous locus. Each circle is a measurement from a single AFD soma normalized to the median at 15°C. Black circles and vertical lines show median and quartiles, respectively. ** and ***: different at P<0.01 and 0.001, respectively compared to values at 15°C within each genotype (one-way ANOVA with Dunnett’s correction). ◊◊◊: different at P<0.001 compared to wild-type within a temperature condition (one-way ANOVA with Dunnett’s correction). 2 independent experiments.

**C)** Quantification of CRH-1 PS nuclear signal intensity in AFD in two independent transgenic lines compared to the unphosphorylatable CRH-1 PS [KID(S48A)] in the indicated temperature conditions. Values are calculated from individual mean projections as the sum nuclear intensity divided by total cell intensity. Each circle is the value from a single AFD neuron. Horizontal and vertical bars indicate mean and SEM, respectively. ** and ***: different at P<0.01 and 0.001, respectively compared to values at 15°C within each genotype (one-way ANOVA with Dunnett’s correction). ◊◊◊: different at P<0.001 compared to wild-type within a temperature condition (one-way ANOVA with Dunnett’s correction). CRH-1 PS Line 1: 8 independent experiments; CRH-1 PS Line 2 and CRH-1 PS [KID(S48A)]: 3 independent experiments.

**D)** Radial cumulative histogram showing temperature-induced changes in CRH-1 PS signal distribution in AFD (see Methods). Data from Figure S2C and Figure 3F. Inset shows magnification of X axis values corresponding to 50% of total signal.

**E)** CRH-1 PS [KID(S48A)] signal distribution in AFD in the indicated temperature conditions as shown by mean projections of nucleus aligned images and mean line scan profiles. Shaded area around the mean line is +/-SEM. n>50 AFD cells per condition; 3 independent experiments.

**Figure S3. Related to Figure 4. RCAN-1 regulates temporal dynamics of *ins-39* expression in AFD.**

**A,E)** (Left) Quantification of fluorescence intensities of the *ins-39* transcriptional reporter in AFD at 15°C in the shown genetic backgrounds. Each circle is a measurement from a single AFD soma. (Right) Quantification of fold-increase of fluorescence intensities of the *ins-39* transcriptional reporter in the indicated temperatures and temperature shift conditions in the shown genetic backgrounds. Each circle is a measurement from a single AFD soma normalized to the median at 15°C. Black circles and vertical lines show median and quartiles, respectively. ***: different at P<0.001 compared to values at 15°C within each genotype (one-way ANOVA with Dunnett’s correction). ◊. ◊◊, ◊◊◊: different at P<0.05, 0.01, and 0.001, respectively. compared to wild-type within a temperature condition (one-way ANOVA with Dunnett’s correction). 2 independent experiments: *cebp-1* and *tax-6 (lof)*); 3 independent experiments: *tax-6(gof)*.

**B)** Raw total fluorescence values of *ins-39::SL2::gfp* reporter in the indicated mutant backgrounds. Each circle is a measurement from a single AFD soma. Black circles and vertical lines show median and quartiles respectively. ***: different at P<0.001 compared to values at 15°C within each genotype (one-way ANOVA with Dunnett’s correction). ◊, ◊◊, ◊◊◊: different at P<0.05, 0.01, and 0.001, respectively. compared to wild-type within a temperature condition (one-way ANOVA with Dunnett’s correction). 10 independent experiments. Alleles used were *mef-2(gv1), rcan-1(tm1925),* and *crh-1(tz2)*.

**C)** Batch effects observed in *ins-39::SL2::gfp* reporter expression in the *rcan-1;mef-2* mutant background. Black circles indicate the mean fold change relative to Day 1. Experiments were performed over three months. Error bars show SEM.

**D)** Mean 2D nuclear fraction value of CRH-1 PS levels in the indicated genotypes and conditions as a function of expression cut off value. Inter-group relationships are not dependent on general sensor expression level. Data from figure 4D. The *rcan-1(tm1925)* allele was used.

**Figure S4. RCAN-1 regulates AFD thermosensory response plasticity.**

GCaMP traces from AFD in adult animals grown under the indicated conditions during a linear temperature ramp at 0.1°C/sec. Thick lines and shading: average ΔF/F change and SEM, respectively. Dashed vertical lines: *T*_AFD_* for wild-type animals. Wild-type *rcan-1* sequences were expressed in AFD under the *gcy-8* promoter as a stably integrated single copy transgene (see Materials and Methods). 3 independent experiments.

**Table S1. Related to all Figures.** Strains used in this work. Excel spreadsheet.

**Table S2. Related to all Figures.** Gene editing reagents used in this work. Excel spreadsheet.

## MATERIALS and METHODS

### *C. elegans* growth and genetics

Animals were grown at 20°C on nematode growth media (NGM) agar plates seeded with *E. coli* OP50. Strains were constructed using standard genetic methods, and genotypes were verified using PCR and/or Sanger sequencing to confirm the presence of molecular changes. Well-fed one-day old adult hermaphrodites were used for all analyses. Strains used in this work are listed in Table S1.

### TRAP-seq

TRAP from AFD was performed essentially as described previously ^15^ with minor modifications. Animals were cultured on 15 cm NGM plates seeded with 2 mL of 10X concentrated *E. coli* HB101. Each plate contained ∼50,000 growth-synchronized one-day old adult animals. A total of 10 plates were used per sample. Animals were grown at 20°C to the L4 larval stage, then shifted to 15°C for 16 hrs. Animals were then harvested directly from 15°C or shifted to 25°C for 1 hr or 4 hrs prior to harvesting. Animals were flash frozen within 15 mins following removal to minimize temperature variation. ∼1000 μg total RNA was used for each immunoprecipitation. Enrichment of AFD-specific transcripts was validated by detection of *gcy-8* mRNA from 3 ng of immunoprecipitated RNA using a OneStep RT-PCR kit (QIAGEN). The affinity matrix used for AFD ribosome purification was prepared using an anti-GFP antibody (RRID:AB_2716737), biotinylated protein L (Thermofisher # 29997), and streptavidin conjugated magnetic beads (Thermofisher # 65601).

Library preparation and sequencing was performed at the MIT BioMicro Center. The Takara SMART-Seqv4 and Illumina Nextera XT kits were used to generate cDNA and the sequencing libraries. Sequencing reads were adapter trimmed using cutadapt with the following options: quality cutoff 20, --trim-n, --minimum-length=50, and then mapped to the *C. elegans* genome (WBcel235/ce11), and counted using STAR with –quantMode GeneCounts.

Differential gene expression analysis was performed in R using DESeq2 ^98^. Differentially expressed genes passing a log2 fold change threshold of 2 were clustered into co-expression modules in Python2 using the DPGP_cluster function (https://github.com/PrincetonUniversity/DP_GP_cluster.git) ^21^. GO analyses and plots were generated using the clusterProfiler, and GOSemSim packages in R.

### CRISPR/Cas9-based gene editing

crRNAs, tracrRNAs and Cas9 protein were obtained from Integrated DNA Technologies (IDT). crRNAs were designed using Benchling’s built in CRISPR design tool. All fusion and *SL2* reporter tags were engineered directly before or after target gene stop codons, respectively.

#### Insertion of reporter or AID sequences

*SL2::H2B::gfp11*, *gfp11,* or *AID* sequences were amplified from gBlock Gene Fragments (IDT) using PCR primers with 5’ overhangs containing ≥ 50bp of DNA homology flanking the Cas9 cut site. *SL2::H2B::gfp11* tags were generated in two steps. First, *SL2::gfp11* sequences were amplified with homology arms from a gBlock and inserted into target gene loci. Next, the *H2B* coding sequence was amplified from plasmid DNA with homology arms designed for in-frame insertion at the *gfp11* N-terminus. The *H2B* coding sequence was inserted into all *SL2::gfp11* intermediate targets using the same crRNA and repair template. *gcy-18::SL2::gfp* was derived from *gcy-18(oy165[gcy-18::gfp])* ^15^ by inserting an *SL2* sequence upstream of *gfp*. Injection mixes for CRISPR/Cas9-mediated gene edits were prepared as described ^99^ and injected into the gonads of one-day old adult animals. A fluorescent co-injection marker (*unc-122p::dsRed*) was added to the injection mix at 50 ng/μl to identify transgenic animals.

#### cebp-1 knockout

Stop cassettes were inserted into *cebp-1* sequences following the protocol for ssDNA repair templates as described ^99^. ssDNA repair templates were obtained from Genewiz as 90nt oligos with standard desalting. Repair templates consisted of a 43 nt STOP-IN cassette ^100^ with 24 nt and 23 nt of homology upstream and downstream, respectively.

#### Generation of ins-39 and gcy-18 alleles with a CRE mutation

Predicted CREs upstream of *gcy-18* and *ins-39* were mutated to contain adenines at every position in the motif. Repair templates for these edits were obtained from Genewiz as 90nt ssDNA oligos with standard desalting. Repair templates contained mutant CRE sequences of variable length flanked by homology arms of maximum size such that the total length did not exceed 90nt.

#### Insertion of single copy transgenes

Single copy *gcy-8*p*::rcan-1* transgenes were stably integrated into the genome via a miniMos transposon ^101^. Other single copy transgenes were inserted into the genome using the CRISPR-assisted knock-in method described previously ^102^. Briefly, *gfp(1-10)* or *TIR1* sequences were cloned into a donor plasmid containing 2638 bp *ttx-1* upstream regulatory sequences and ∼1600 bp homology arms targeting a region of chromosome II. The donor plasmid also encoded hygromycin resistance and the *sqt-1(e1350)* selection markers as well as an *hsp-16.41* promoter-driven *Cre* recombinase. Selection marker and *hsp-16.41*p::*Cre* coding sequences were flanked by *loxP* sites to allow excision of selection markers by heat shock after integration. Injection mix was prepared as outlined ^102^. The donor plasmid, Cas9/sgRNA-encoding plasmid, and a *myo-2*p*::mCherry* co-injection marker were injected together into the gonads of one-day old adult hermaphrodites. Stable integration was identified via hygromycin resistance, roller phenotype, and absence of *myo-2*p*::mCherry* expression. Individual animals were cloned out and allowed to reproduce. Progeny were heat-shocked at 34°C for 4 hrs to activate Cre expression and excise selection markers. Reagents used for gene editing are listed in Table S2.

### CRH-1 phosphorylation sensor

The CRH-1 PS sensor plasmid (KID::NES::SL2::KIX::H2B) was made in three sequential steps via Gibson assembly using the NEB HiFi Gibson assembly master mix. First, a CRH-1 KID::NES sequence synthesized as a gBlock (IDT) was inserted downstream of the *ttx-1* promoter and in-frame with *gfp* coding sequences into an expression vector. Next, a SL2::KIX gBlock (IDT) was inserted into this construct. Finally, KID::NES::SL2::KIX and H2B encoding sequences were PCR-amplified with overlapping homology and assembled to generate the final sensor plasmid in which H2B was fused in frame downstream of KIX. The S48A mutation was generated via site-directed mutagenesis (NEB). Sensor plasmids were injected at 5-15 ng/μl with an *unc-122*p*::dsRed* co-injection marker at 50 ng/μl.

### Microscopy

#### Fluorescent reporters

To quantify expression of fluorescent reporters, L4 larvae were picked from 20°C and transferred to a 15°C incubator for >6 hrs, following which one set was moved to 25°C overnight. On the following day, animals were shifted from 15°C to 25°C for 1 hr or 4 hrs. Animals were then immobilized with 10 mM tetramisole, mounted on 10% agarose pads on slides, and imaged on DM6000 Leica inverted microscope with a CSU-W1 spinning disk confocal unit and a ZL41 sCMOS camera using either a 63x or 100x oil objective and *z*-step sizes of 0.25μM. 3D volumes were captured for each animal using the Fusion software from Oxford Instruments. Images were processed in MATLAB using custom scripts to draw and select ROIs by intensity thresholding. Expression was quantified from a maximum projection as corrected total cell fluorescence (CTCF) using the following equation:

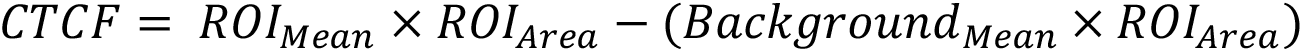

#### CRH-1 phosphorylation sensor

Images of the CRH-1 phosphorylation sensor were quantified using a custom MATLAB image analysis pipeline. Cells were first segmented in three dimensions and embedded in 3D NaN arrays to exclude background during projection. Each cell volume was then manually rotated to align its longest axis with the x-axis and to maximize visibility of nuclear/cytoplasmic contrast. Mean intensity projections were generated from the rotated volumes and individually min/max normalized. Nuclear centers were manually annotated on each projection, and these annotations were used to align all projections into a 3D volume centered on the nucleus. The aligned projections were averaged to generate the images shown in Figures 3F, 4C and S2E. In addition, these images were cropped to retain pixels with a minimum of 50% coverage and a Gaussian blur was applied to blend elements of the individual nuclear aligned images.

To quantify the fraction of nuclear signal, a circular region of interest with a manually adjusted radius was drawn around the nucleus in each mean projection. The nuclear fraction was calculated as the sum fluorescence intensity within the nuclear region divided by the total fluorescence intensity of the cell on the 2D projection. Line scan profiles were generated from fluorescence intensity values sampled along a straight line parallel with the cells longest dimension and through the manually annotated nuclear centers.

To generate the radial cumulative histogram in Figure S2D, a circle was drawn starting at the nuclear center of individual mean projections and expanded by 1 pixel/step until the entire cell was contained in the circle. The sum intensity within the circle at each step was divided by the total cell intensity to give the fraction of intensity contained within each circle.

### Auxin-induced degradation

Synthetic auxin (Sigma-Aldrich #317918) dissolved in 70% ethanol was diluted 1:100 in NGM to obtain a final concentration of 4 mM on the growth plate. Control plates contained 70% ethanol alone. Auxin and ethanol-containing NGM plates were seeded with *E. coli* OP50. 10-15 animals were placed on individual plates and subjected to the indicated auxin exposure and temperature experience prior to imaging.

### Calcium imaging

Calcium imaging was performed using one-day old hermaphrodites expressing GCaMP6s in AFD ^91^. Following the indicated temperature shifts, animals were immobilized in 10mM tetramisole and mounted on agarose pads on a glass coverslip and transferred to the temperature-controlled stage of an upright Zeiss Axiskop2 Plus microscope. This microscope stage is equipped with a thermoelectric cooler (TEC) bonded to a liquid-cooled heat sink for precise temperature control. The TEC delivered a linear temperature ramp rising at 0.1°C/s. Recordings of calcium dynamics were made using Metamorph and analyzed in MATLAB using custom scripts (https://github.com/SenguptaLab/AFD_gene_expression.git).

## Statistical analyses

All results shown are from at least two biologically independent experiments; the number of experiments is indicated in each figure legend. TRAP-seq analyses was performed with data from four independent experiments. Statistical tests for reporter quantifications were performed in MATLAB using built-in functions for one-way ANOVA and Dunnett’s correction. TRAP-seq statistical analyses were performed in DESeq2 following the pipeline for tissue and batch correction for time series expression data described previously ^103^.

## Data and code availability

Data for all figures are available on FigShare (https://figshare.com/s/0c23d4b56dab4f85dc29). TRAP-Seq data have been deposited to GEO (Bioproject PRJNA1503593). MATLAB image analysis code used for CRH-1 PS and reporter quantification is available at https://github.com/SenguptaLab/Bates2026.git.

## Acknowledgements

We are grateful to the *Caenorhabditis* Genetics Center for strains, Jihye Yeon for critical comments on the manuscript, and the Sengupta lab for advice and input. This work was supported in part by the NIH (R35 GM122463 – P.S., and T32 GM139798 and F31 NS134251 – S.B.).

