## Supplementary figures and images for "A repressive regulatory cascade shapes temporal patterning of activity-regulated gene expression in a defined sensory neuron type"

### Supplement

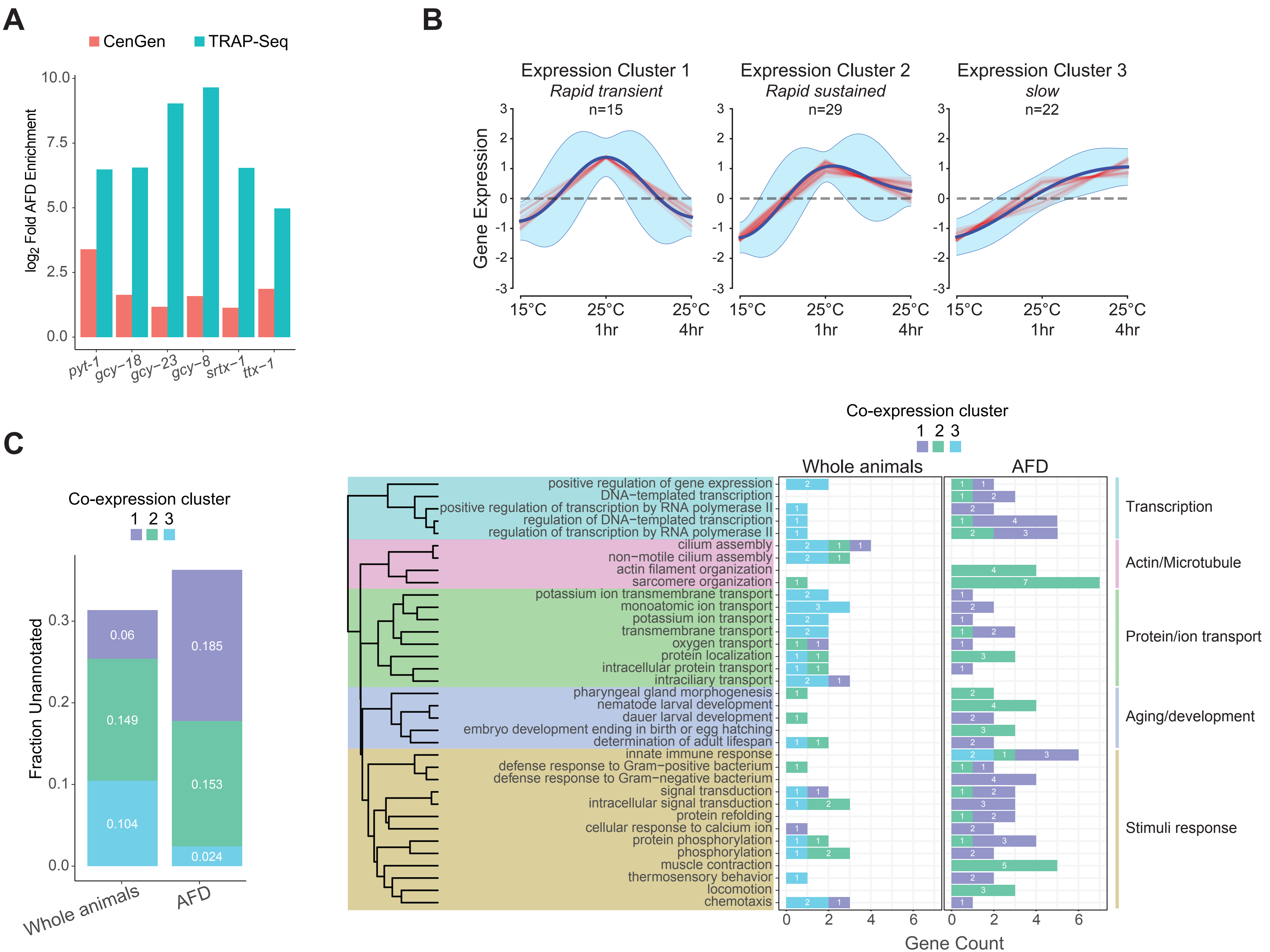


**Bates et al., Figure S1**

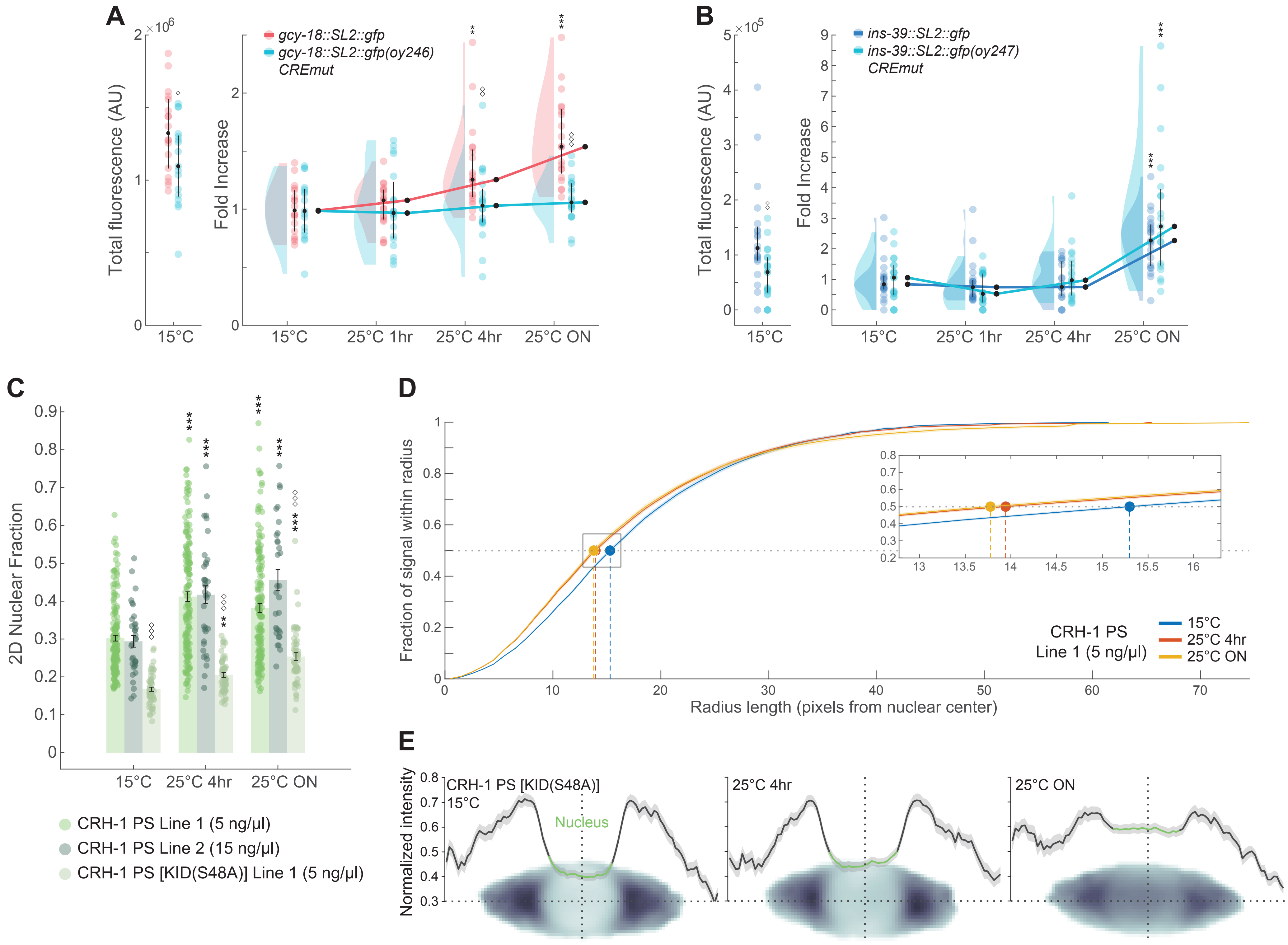


**Bates et al., Figure S2**

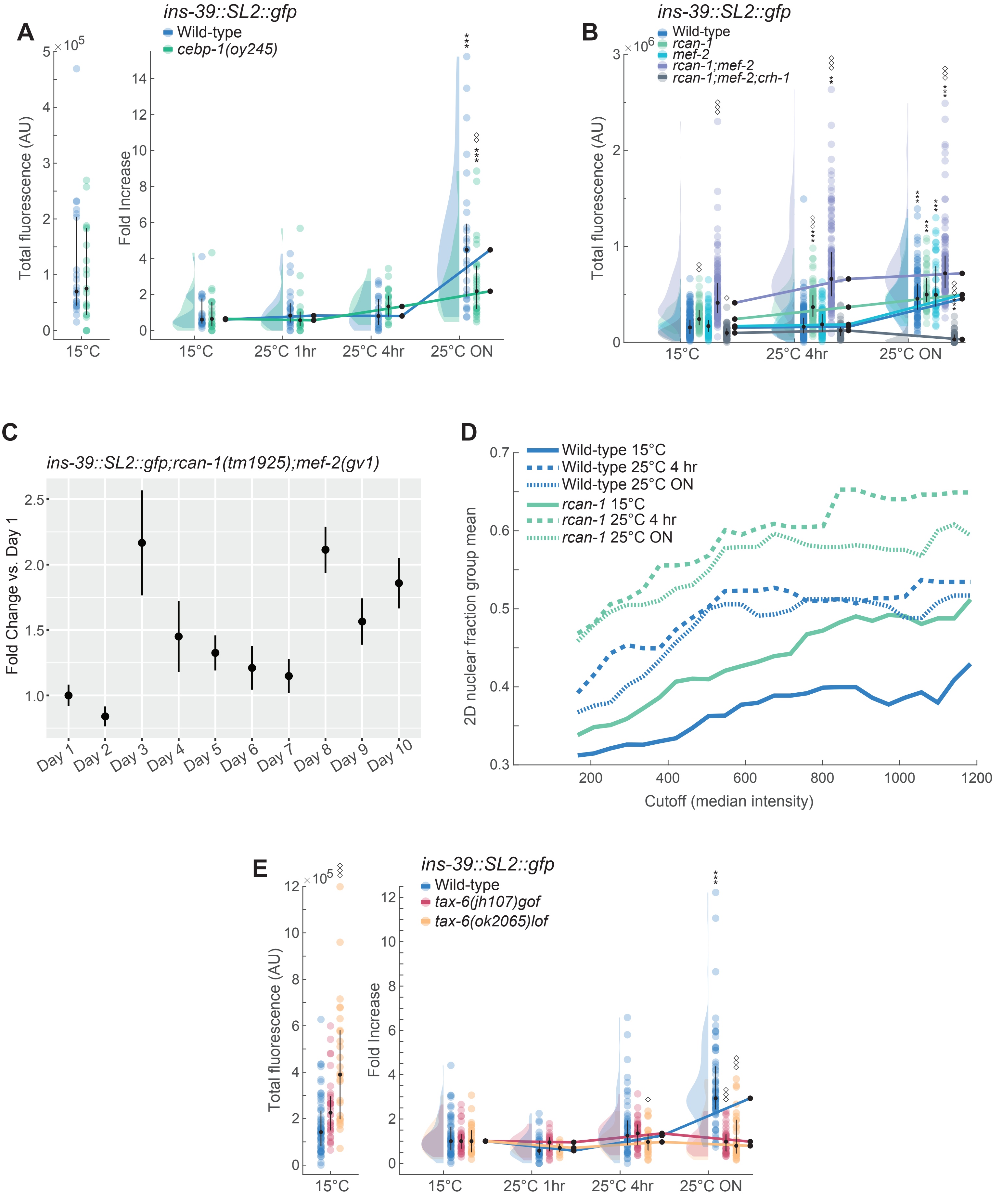


**Bates et al., Figure S3**

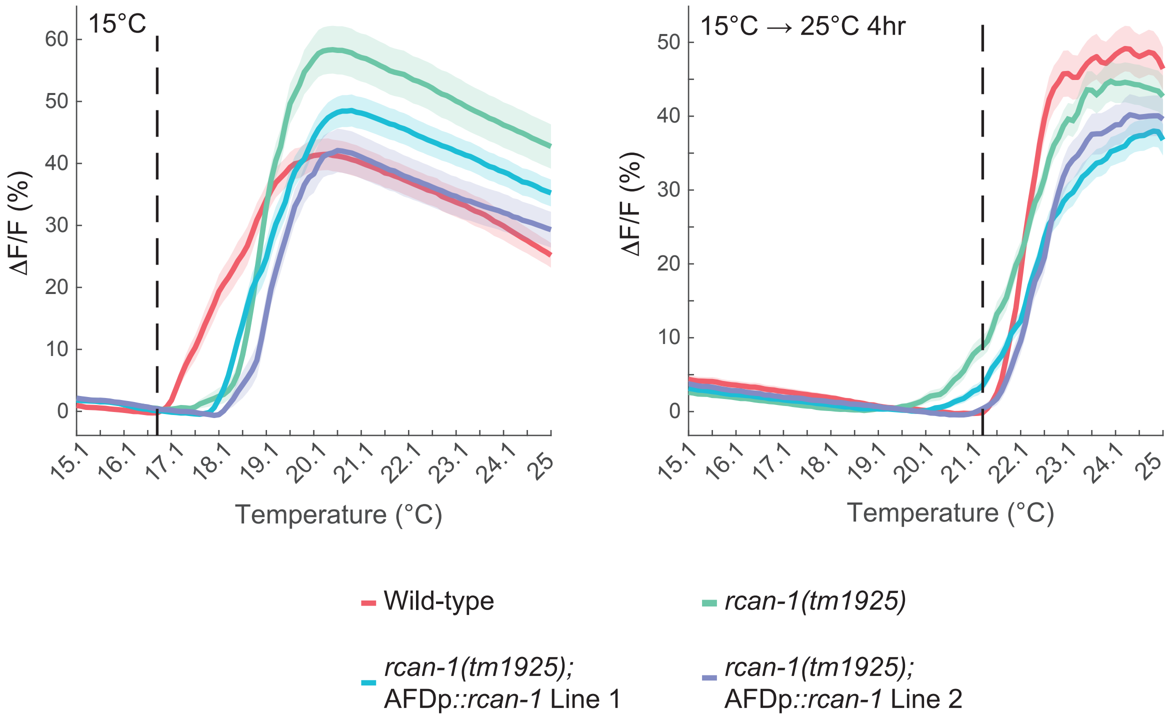


**Bates et al., Figure S4**
